# DeLoop: a deep learning model for chromatin loop prediction from sparse ATAC-seq data

**DOI:** 10.1101/2023.11.01.564594

**Authors:** Yihai Luo, Zhihua Zhang

## Abstract

Deciphering gene regulation and understanding the functional implications of disease-associated non-coding variants require the identification of cell-type-specific 3D chromatin interactions. Current chromosome conformation capture technologies fall short in resolution when handling limited cell inputs. To address this limitation, we introduce DeLoop, a deep learning model designed to predict CTCF-mediated chromatin loops from sparse ATAC-seq data by leveraging multitask learning techniques and attention mechanisms. Our model utilizes ATAC-seq data and DNA sequence features, showcasing superior performance compared to existing state-of-the-art models, particularly under low read depth conditions, enabling accurate chromatin loop inference when sufficient cells are infeasible. In addition, generalizing across cell types, DeLoop proves effective in de novo prediction tasks and its potential for predicting functional interactions.

## Main

The genetic information of the human genome is stored in the linear DNA, while the folding of chromosomes forms the complex three-dimensional structure of the genome. This three-dimensional genome organization, particularly the formation of chromatin loops, is crucial for gene expression regulation^1^. As a fundamental architectural feature of the 3D genome, the chromatin loop involves the physical interaction and spatial proximity of distant genomic regions, often separated by hundreds of thousands to millions of base pairs along the linear DNA sequence. These loops are facilitated by the bending and folding of the chromatin fiber, allowing specific genomic elements, such as enhancers, promoters, and other regulatory sequences, to come into close contact. Key transcription factors like CCCTC-binding factor (CTCF) dynamically regulate genes through these loops, influencing cellular function, morphology, and fate decisions. The chromatin loops exhibit considerable heterogeneity in different cell lines and within the same cell during different cell cycles^2^. These cell-specific chromatin loops contribute to cellular heterogeneity, impacting cell fate and function. Therefore, detecting chromatin loops is essential in studying various aspects of the genome, including its three-dimensional structure, cell differentiation, and the mechanisms of disease occurrence^3^.

Genome-wide chromosome conformation capture techniques such as Hi-C^4^ and ChIA-PET^5^ have been devised to capture the intricate three-dimensional conformation of chromosomes, revealing chromatin interactions. Advances like Hi-ChIP have improved the sensitivity and efficiency of capturing chromatin interactions and reduced sequencing costs^6^. Nonetheless, data resolution from techniques like Hi-C and ChIA-PET depends on sequencing depth, requiring substantial financial, temporal, and human resources. This challenge becomes pronounced when investigating a limited number of cells where obtaining a reliable contact map is unattainable due to insufficient read depth. Therefore, computational methods for predicting chromatin loops are crucial for studying the 3D structure of the genome.

Predictive methods fall into two categories: those relying solely on sequence features (e.g., CTCF-MP^7^ and ChiNN^8^) and those integrating multi-omics data (e.g., Lollipop^9^ and CISD-loop^10^). Sequence-based models, though not influenced by sequencing depth, struggle with cross-cell line predictions and fail to capture cell line heterogeneity. Multi-omics models demonstrate superior performance but require high-quality input data, posing challenges in obtaining multiple omics data from the same sample. This prompts the exploration of methods minimizing input signals to achieve precise loop prediction, addressing a crucial question in the study of three-dimensional genome structure. In addition, to our best knowledge, there is no multi-omics approach optimized for sparse epigenetic signals, limiting 3D genome research in low read-depth samples.

In this study, we introduce DeLoop, a deep learning model by leveraging multitask learning techniques and attention mechanisms to predict CTCF-media chromatin loops from sparse ATAC-seq data and DNA sequence features. DeLoop outperforms state-of-the-art deep learning models designed for similar interaction prediction tasks and is robust to the background noises and sparsity from low read depth ATAC-seq data.

## Result

### 1 DeLoop is a deep-learning model that predicts chromatin loops from ATAC-seq data

Based on our previous model CISD-loop, it has been demonstrated that chromatin accessibility can be leveraged to predict chromatin interactions^10^. Given the widespread availability and cell-type specificity of chromatin accessibility information derived from ATAC-seq, we have incorporated ATAC-seq signals and DNA sequences as inputs for our model. In addition, Agarwal et al. emphasized the significance of anchor distance in chromatin loop prediction^11^. Therefore, we developed a novel deep-learning multimodal model, DeLoop, integrating DNA sequence features and chromatin accessibility information to predict CTCF-media chromatin loops. Specifically, DeLoop takes two 2048bp one-hot encoded DNA sequences of loop anchors, their corresponding chromatin accessibility scores derived from ATAC-seq data, and the loop distance to predict the probability of whether these two anchors may form a chromatin loop.

We attend positive loops from the CTCF ChiA-PET dataset and only use the loop anchors under 5kb long since chromatin loop anchors can be down to 1kb in size. The low-resolution interactions could lead to the introduction of noise to the training dataset. The corresponding distance-matched negative datasets were generated approximately five times as the positive sample from three sources, including pairs of non-directly or indirectly connected loop anchors, random pairs of peaks from CTCF ChIP-seq data, and random pairs of peaks from ATAC-seq data. To simulate sparse ATAC-seq data, we initially downsampled the raw ATAC-seq reads to 40 million reads, creating a standard ATAC-seq dataset as a reference. Subsequently, we downsampled this standard dataset to 1 million reads, generating sparse ATAC-seq data. Throughout the training phase, we employed the sparse ATAC-seq data for data augmentation, enhancing the model’s performance in handling sparse data. This strategy was implemented to address sparsity challenges and bolster the model’s robustness in scenarios where only low read depth ATAC-seq data is available.

ATAC-seq signals and DNA sequences are passed through two DenseNet-based feature extractors in the DeLoop architecture (**Fig. 1**). A transformer-based integration module is used to fuse features from two modalities to anchor features, allowing cross-modal information exchange. This integration module generates representations of the anchor features for two candidate loop anchors, subsequently processed through an additional integration module to fuse the features of both anchors into a coherent loop representation. Finally, this loop representation and anchor distance are input into a multi-layer perceptron classifier to predict chromatin interactions.

**Figure 1.**
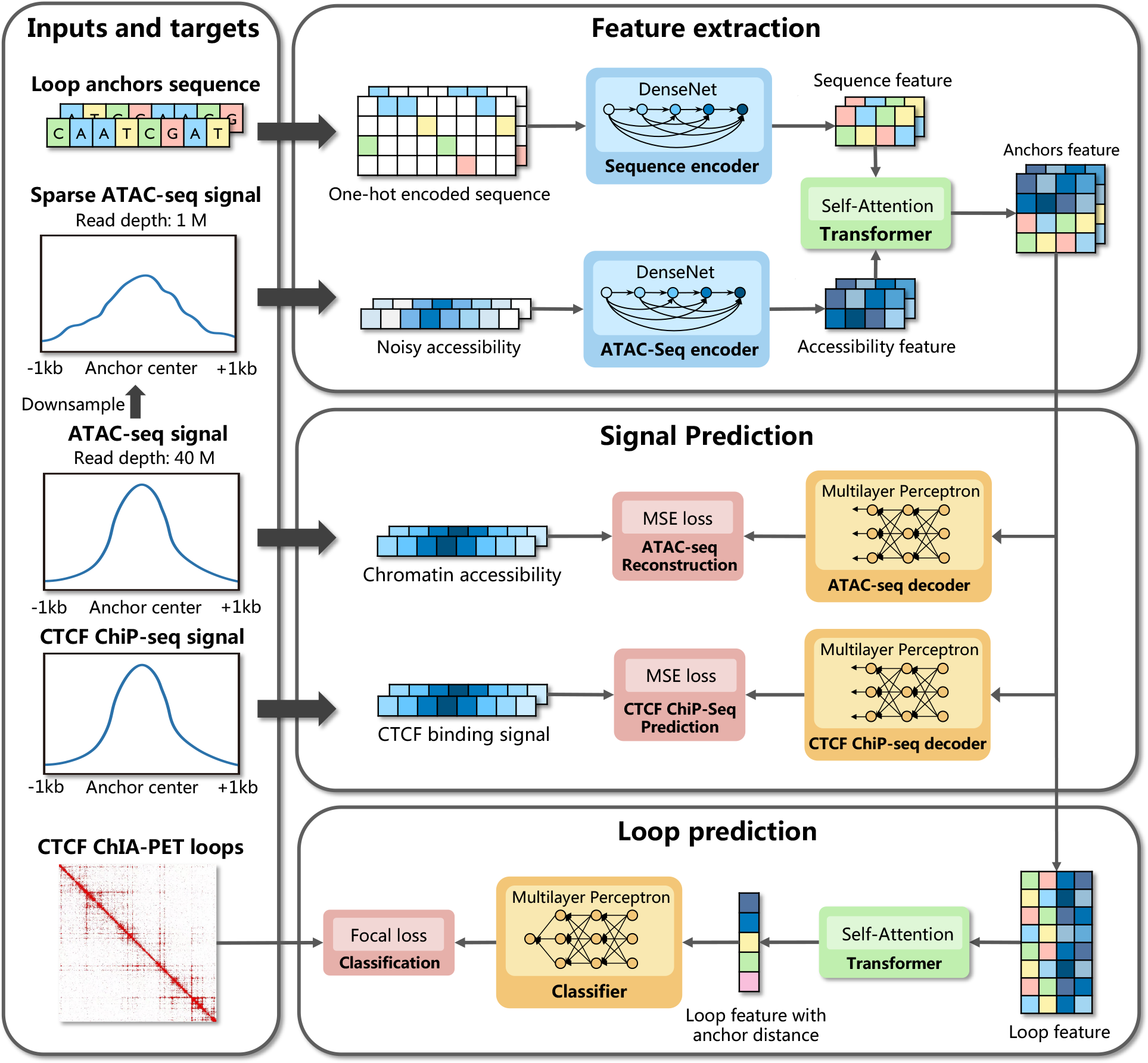
The pipeline and model architecture of DeLoop.

In order to optimize the capability of the cross-modal integration module in seamlessly integrating features from DNA sequence and ATAC-seq signal, we introduced lightweight ATAC-seq signal and CTCF ChiP-seq signal decoders following the anchor features. Since the sparse ATAC-seq signals are used for data augmentation, by learning to reconstruct the sparse ATAC-seq signals to standard ATAC-seq signals, the model is incentivized to utilize sequence features for the denoising of ATAC-seq signals while capturing CTCF binding information at the same time (**Methods**). Adding additional loss functions contributes to a refined precision in chromatin loop classification. Notably, DeLoop attains the highest average precision when employing a multi-task loss involving loop classification, ATAC-seq signal prediction, and CTCF binding signal prediction.

This result demonstrates that features learned from epigenomic signal prediction can be transferred to loop prediction (**Table 1**).

**Table 1.** Multitask loss improves DeLoop’s performance in both standard and sparse ATAC-seq data.

| Experiment | Standard ATAC-seq data |  |  | Sparse ATAC-seq data |  |  |
| --- | --- | --- | --- | --- | --- | --- |
|  | Precision | Recall | AP | Precision | Recall | AP |
| $Loss_{cls}$ | 0.6696 | 0.9426 | 0.9067 | 0.6251 | 0.9198 | 0.8695 |
| $Loss_{ATAC} + Loss_{cls}$ | 0.6817 | 0.9463 | 0.9098 | **0.6710** | 0.9004 | 0.8743 |
| $Loss_{CTCF} + Loss_{cls}$ | 0.6797 | **0.9480** | 0.9121 | 0.6673 | 0.9090 | 0.8821 |
| $Loss_{CTCF} + Loss_{ATAC} + Loss_{cls}$ | **0.7072** | 0.9415 | **0.9148** | 0.6656 | **0.9254** | **0.8824** |

To assess the predictive capacity of ATAC-seq signals and loop anchor distance information for chromatin loops, we trained DeLoop using various combinations of these features and sequence features. Subsequently, we conducted testing on diverse ATAC-seq datasets with varying read depths. Remarkably, the model trained with ATAC-seq signals and loop anchor distance information exhibited superior performance compared to the model relying solely on sequence features. The model incorporating all three features demonstrated the highest performance across standard and sparse ATAC-seq data, as illustrated in **Fig. 2a, b**. For further evaluation, we defined all candidate loop anchors overlapping with real loop anchors as active ones. Forming a loop between two candidate anchors required both to be active interacting anchors. Without supervised training specifically on active loop anchors, the anchor features effectively separate active and non-active anchors during the training process (**Fig. 2c**). To investigate how sequence and accessibility information play a role in distinguishing active and non-active loop anchors, we performed Linear Discriminant Analysis (LDA) dimensionality reduction on sequence, accessibility, and anchor features for the test dataset (**Fig. 2d,e,f**). Notably, while using sequence and ATAC-seq signals in isolation could separate active and non-active loop anchors, the anchor features integrating both elements achieved the highest class separation. The result demonstrates that sequence and ATAC-seq signals contain complementary information for chromatin loop prediction, and DeLoop’s encoders can capture necessary information for chromatin loop prediction. This not only highlights the importance of combining both features for predicting chromatin loops but also validates the effectiveness of our encoders and integration module in successfully extracting and merging these two features.

**Figure 2.**
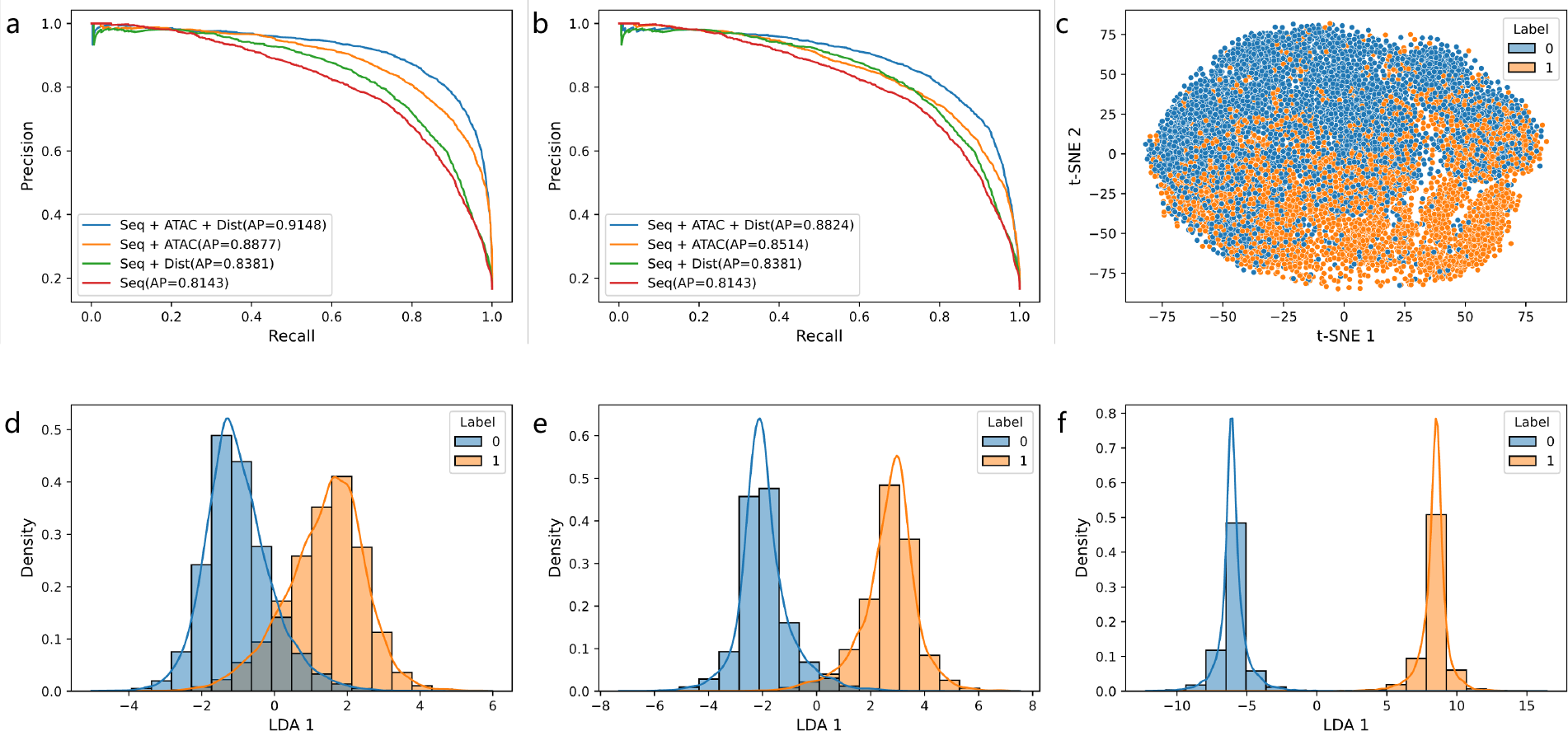
Anchor distance and accessibility information improve loop prediction. **a**. Precision-recall curves of deLoop in standard ATAC-seq data with a read depth of 40 million reads. **b**. Precision-recall curves of deLoop in standard ATAC-seq data with a read depth of 1 million reads. **c**. t-SNE dimensionality reduction on anchors feature. **d-e**. Dimensionality reduction using LDA on features from sequences, ATAC-seq, and anchors.

### 2 ATAC-seq features Multitask learning enables loop prediction in sparse ATAC-seq data

Many deep-learning models have been developed to predict chromatin interactions. To evaluate the performance of DeLoop, we select the state-of-the-art models DeePPHiC and ChiNN to compare with DeLoop. For a fair comparison, We trained and tested these models on the same dataset from three human cell lines (GM12878, K562, HelaS3) separately. DeLoop outperformed the other two models in predicting chromatin loops from standard and sparse ATAC-seq data (**Fig. 3 a,b,c**).

**Figure 3.**
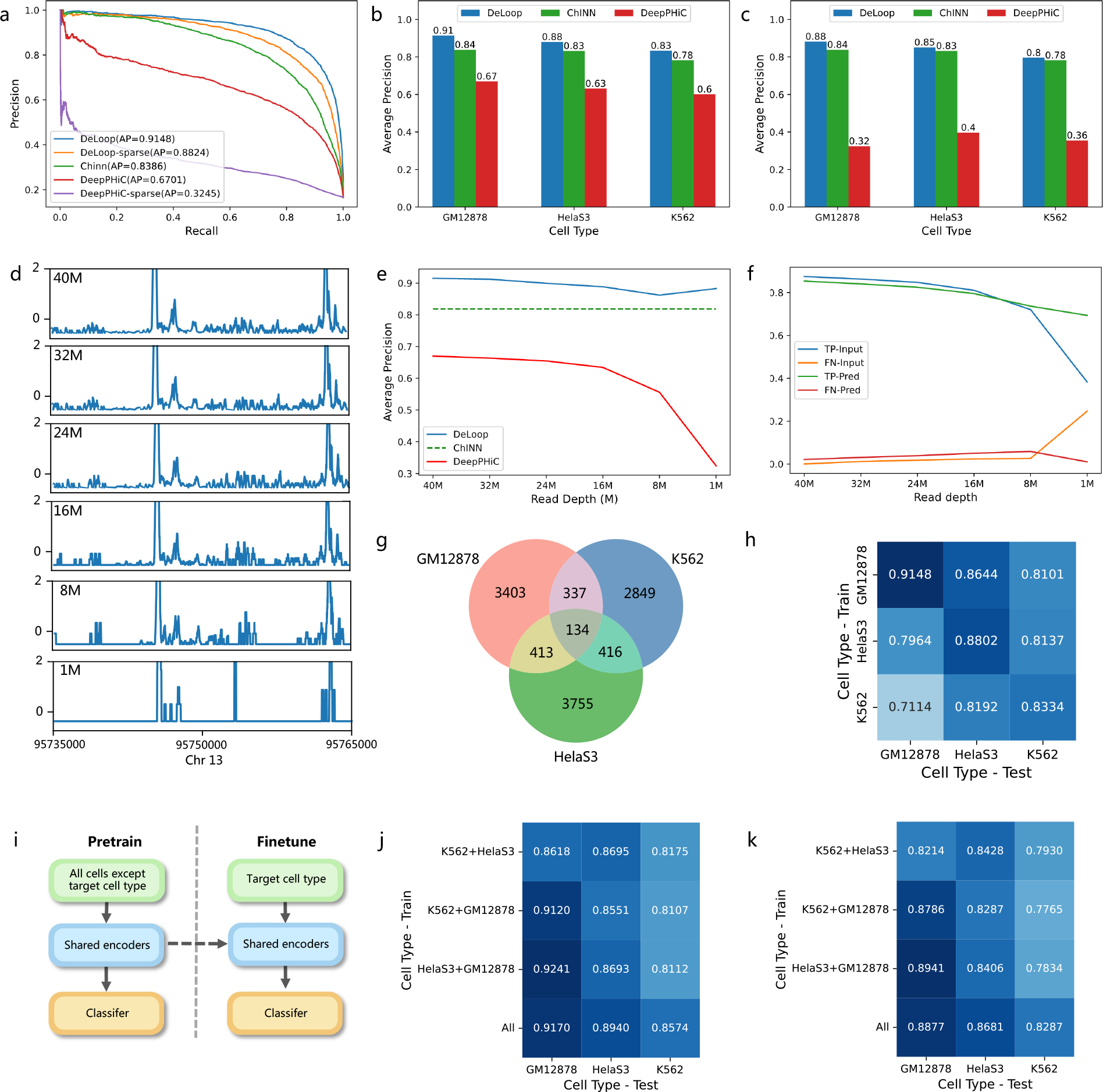
DeLoop is the state-of-the-art CTCF-media loop prediction model. **a**. Performance comparison of DeLoop, DeepPHiC, and ChiNN in GM12878. **b**. Average precision in standard ATAC-seq data. **c**. Average precision in sparse ATAC-seq data. **d**. Normalized accessibility signal across various ATAC-seq data read depths. **e**. Model performance across ATAC-seq data with varying read depths. Notably, ChiNN, relying solely on DNA sequence and anchor distance, remains unaffected by ATAC-seq data sparsity. **f**. The ratio of true positive peaks and false negative peaks under different read depths. **g**. Venn plot depicting chromatin loop intersections in the test dataset across three human cell lines. **h**. Average precision of deLoop in cross-cell prediction task. **i**. Transfer learning paradigm employed by DeLoop. **j**. Average precision of DeLoop using transfer learning with standard ATAC-seq data. **k**. Average precision of deLoop using transfer learning using sparse ATAC-seq data.

Since the ChiNN only makes use of DNA sequence and distance information, we replace the ATAC-seq signal with all zero signals in our trained model to evaluate DeLoop’s performance when accessibility information is not available. Without specific training on data with all zero signals, DeLoop achieved an average precision of 0.8542 in GM12878, surpassing ChiNN’s 0.8186. These results demonstrate that DeLoop’s design is more effective than ChiNN in predicting chromatin loops even without ATAC-seq signals.

To further evaluate how sparsity in ATAC-seq data could affect the performance of DeLoop, we test DeLoop on different read depths. We found that, compared to DeepPHiC, whose performance drops dramatically as the read depth decreases, DeLoop exhibits only a marginal drop in performance. This result demonstrates that DeLoop is more robust to sparse ATAC-seq data than DeepPHiC. Analysis of **Fig.3 f** reveals a diminishing ratio of true positive ATAC-seq peaks alongside an increase in false negative and false positive peaks as read depth decreases, particularly from 8 million to 1 million reads. **Fig.3 d** shows the normalized ATAC-seq signals from different read-depth data. Comparing signals from 1 million reads to deeper read depth, positive peaks reduce as the data gets sparser, and a false positive peak can be observed. This result shows that the sparsity in ATAC-seq data could lead to the loss of necessary information for chromatin loop prediction as well as the introduction of background noises, elucidating why DeepPHiC’s performance decreases significantly with reduced read depth.

Since DeLoop makes use of the sequence feature to update the accessibility feature in the cross-modal integration module and learn to denoise and reconstruct sparse ATAC-seq signal, the ratio of true positive ATAC-seq peaks in the ATAC-seq signal that DeLoop prediction decreases more slowly compared to the input ATAC-seq signal as we expected (**Fig.3 f, Supplementary Fig.1**). This result affirms that DeLoop can effectively denoise the sparse ATAC-seq signal and capture the necessary information for chromatin loop prediction since DeLoop makes use of the sequence feature to update accessibility feature in the cross-modal integration module and learn to denoise and reconstruct sparse ATAC-seq signal.

To assess DeLoop’s performance in cell-type-specific de novo prediction tasks, we trained DeLoop on one cell type and tested it on others. As shown in **Fig.3 g**, most loops in the test dataset are cell-type specific. DeLoop can generalize across cell types and achieve acceptable performance in predicting chromatin loops in different cell types (**Fig.3 h**). Incorporating transfer learning further enhances DeLoop’s performance by leveraging features learned from other cells and applying them to the target cell (**Fig.3 i**). Compared to deLoop without transfer learning, pretraining with data from other cell types yields superior performance (**Fig.3 j,k**).

## Discussion

In this study, we developed DeLoop, a deep-learning model that excels in predicting chromatin loops from sparse ATAC-seq data. By using an attention mechanism to incorporate DNA sequence features and chromatin accessibility information, DeLoop surpasses existing models in accuracy and robustness. The architectural framework of DeLoop is characterized by DenseNet-based feature extractors and a transformer-based integration module, highlighting its efficacy in facilitating cross-modal information exchange.

The utilization of multitask learning, including loop classification, ATAC-seq signal prediction, and CTCF binding signal prediction, further refines the precision of chromatin loop classification. Notably, DeLoop’s ability to denoise sparse ATAC-seq signals while capturing essential CTCF binding information sets it apart from other models. Introducing supervision on decoders for ATAC-seq and CTCF ChIP-seq signals enhances the model’s capability to extract meaningful features.

Our comprehensive evaluation demonstrates that DeLoop outperforms state-of-the-art models such as DeepPHiC and ChiNN, even in the absence of ATAC-seq signals. The robustness of DeLoop to sparse ATAC-seq data is a notable advantage, as evidenced by its performance across varying read depths. The model’s ability to generalize across cell types further solidifies its utility in cell-type-specific de novo prediction tasks.

In conclusion, DeLoop represents a significant advancement in chromatin interaction prediction. Its integration of sequence features and epigenomic signals, coupled with its robust performance in sparse data scenarios, positions it as a valuable tool for unraveling the complexities of chromatin loops and their impact on cellular function and fate decisions. Anticipating the expanding availability of single-cell data and potential future upgrades to DeLoop, it is envisioned that this model will prove instrumental in dissecting the 3D genome, particularly when applied to single-cell loop prediction.

## Code availability

The source code will be available after the peer-review process is completed.

## Method

### DeLoop collection and preprocess

We used CTCF ChIA-PET, ATAC-seq, and CTCF ChIP-seq data from three human cell lines (**Supplementary Table 1**). Processed CTCF binding peaks are acquired from the CTCF ChiP-seq dataset. Chromatin loops called from CTCF ChIA-PET are acquired from ChINN (GM12878, HelaS3) and ENCODE (K562). We adopted ChiNN’s pipeline and source code in preparing the chromatin loops dataset.

Chromatin interactions overlapping with ENCODE blacklisted regions at either anchor were excluded. Anchors of retained interactions in each dataset were merged with a 500 bp permissible gap. The resulting clusters were filtered against ATAC-seq peaks, yielding a set of positive samples for each dataset. Distance-matched negative datasets, comprising approximately five times the positive samples, were derived from three sources: pairs of loop anchors without direct or indirect connections, random pairs extracted from CTCF ChIP-seq data, and random pairs sourced from ATAC-seq data. Raw ATAC-seq data is randomly downsampled to 40 million reads as standard ATAC-seq data reference. ATAC-seq data from different read depths is further downsampled from this ATAC-seq data reference. Sampled ATAC-seq data is processed with ENCODE ATAC-seq pipeline with all default parameters^12^.

### DeLoop model training

We employed a multi-task learning loss during the training process to enable the model to perform accurate loop classification while concurrently predicting denoised ATAC-seq data and CTCF binding signals based on the input sparse ATAC-seq data and sequence information. This multi-task loss allows for the transfer of features extracted from the prediction of ATAC-seq and CTCF binding signals to the classification task, thereby enhancing the accuracy of the classification task.

We implement the focal loss^13^ in our classification task defined as:

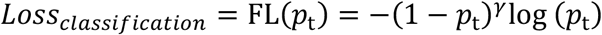

γ is a tunable parameter set to 1 as default .*p*_*t*_ in the above is defined as

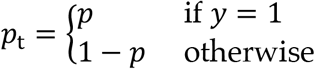

*y* ∈ {0,1} specifies the ground-truth class, and *p* ∈ [0, 1] is the model’s estimated probability for the class with the label *y* = 1. By adding a factory (1 − *p*_t_)^γ^ to the standard cross entropy criterion −log (*p*_t_), the loss function down-weights easy examples and thus focus training on hard negatives.

We use the average mean squared error between predicted and actual signals from two anchors as the loss function for accessibility and CTCF binding prediction. The predicted signal length is set to 64 with a bin size of 32 bp.

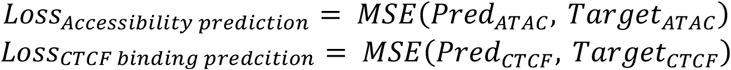

The multitask loss is defined as

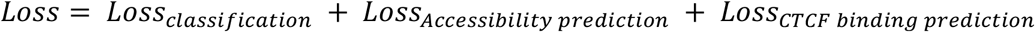

Samples on chromosomes 1-21 were used for training, and samples on chromosomes 4,7,8, and 11 were used for testing, while the rest were used for training. In the training process, to mitigate bias, a resampling approach is employed to ensure a balanced number of positive and negative samples. In the training phase, 30% of ATAC-seq input is randomly downsampled from standard ATAC-seq data to sparse ATAC-seq data.

### DeLoop architecture

DeLoop is implemented with the PyTorch framework. DeLoop consists of two DenseNet-based feature encoders, two multi-layer perception decoders, two transformer-based integration modules, and a multi-layer classifier. DeLoop task two four channels one-hot encoded DNA sequence with the length of 2048bp and two 1-channel accessibility signals attained from ATAC-seq data. For efficiency and robustness, the length accessibility signals and CTCF binding scores predicted by DeLoop is 64 with a bin size of 32bp.

DeLoop’s encoders are developed based on DenseNet^14^ due to its proven ability to encourage feature reuse by dense skip connections, thereby reducing the number of parameters. These connections link preceding layers to all subsequent layers through feature concatenation. This design ensures that each layer directly accesses output gradients during backpropagation, leading to faster network convergence and improved regularization.

We implement the transformer module to introduce the self-attention mechanism, effectively capturing long-range dependencies within sequences and overcoming limitations associated with traditional recurrent neural networks when dealing with lengthy input sequences^15^. Its key components include multi-head self-attention layers and feedforward neural network layers, allowing the transformer to learn and capture intricate structures and patterns within input sequences. In the first transformer-based integration module, features from ATAC-seq and DNA sequences are concatenated to form anchor features. The fusion of features across modalities is achieved through attention mechanisms within the transformer. In the second integration module, features from two anchors are combined into an elongated sequence, enabling the capture of correlations between anchors.

**Supplementary Figure 1.**
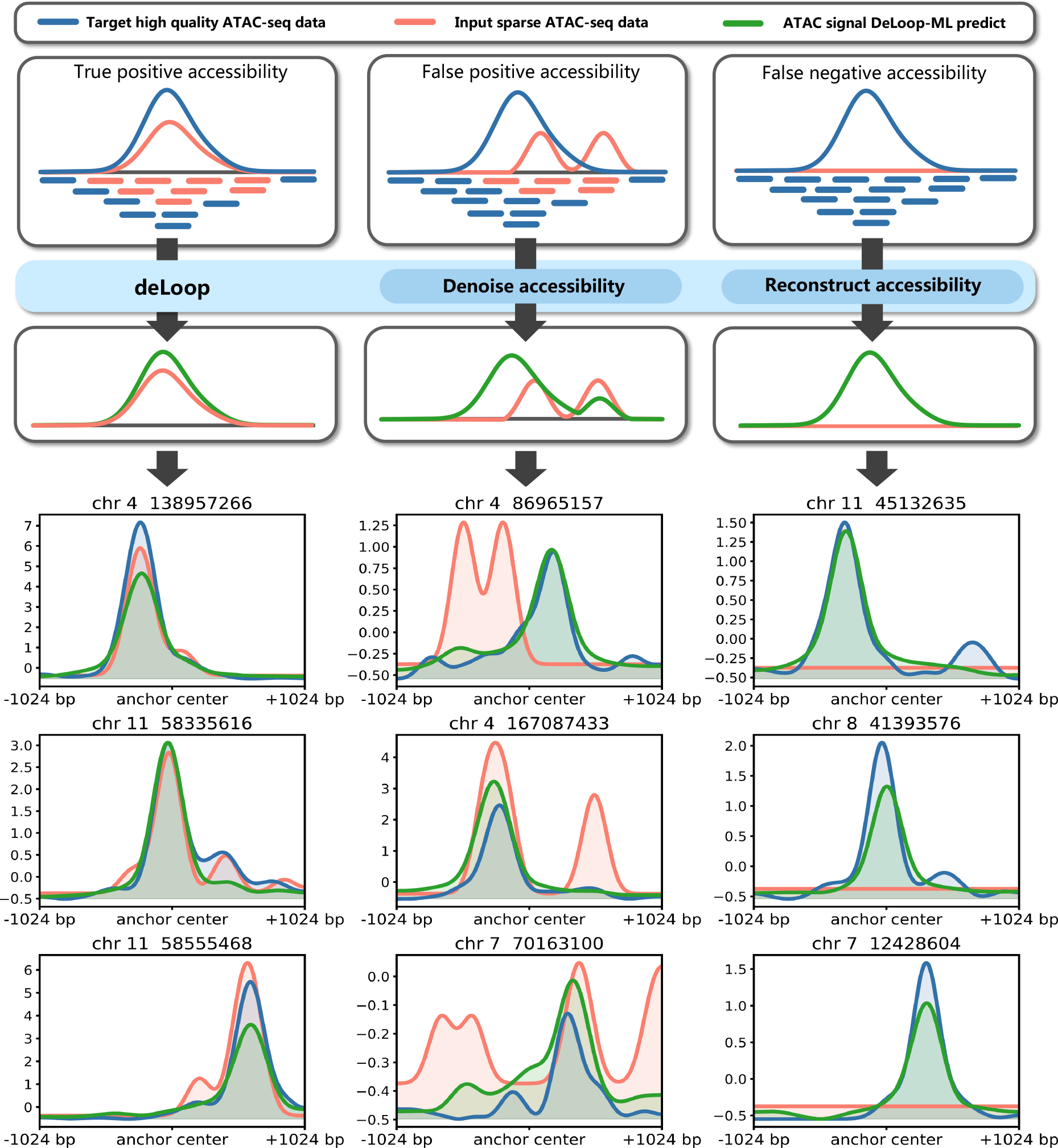
DeLoop improves its performance in sparse ATAC-seq data by accessibility signal denoising and reconstruction. The figure illustrates potential biases introduced in chromatin accessibility information from extremely sparse ATAC-seq data with low library complexity and how DeLoop manages to eliminate these biases. The left column represents the scenario without bias. The middle column depicts situations where signal peaks are offset, or background noise is misinterpreted as signal peaks, resulting in false-positive accessibility. DeLoop demonstrates effective denoising in such cases. The rightmost column illustrates instances where signal peaks cannot be detected due to excessive sparsity, leading to false negatives. DeLoop excels in predicting the required accessibility information based on sequence information.

